# Improving prediction and assessment of global wildfires using neural networks

**DOI:** 10.1101/2020.08.07.241638

**Authors:** Jaideep Joshi, Raman Sukumar

**Author notes:** correspondence may be addressed to Jaideep Joshi or Raman Sukumar.

## Abstract

Fires determine vegetation patterns, impact human societies, and provide complex feedbacks into the global climate system. Empirical and process-based models differ in their scale and mechanistic assumptions, giving divergent predictions of fire drivers and extent. Especially, the role of anthropogenic drivers remains less understood. Taking a data-driven approach, we use an artificial neural network to learn region-specific relationships between fire and its socio-environmental drivers across the globe. As a result, our models achieve higher predictability than previously reported, with global spatial correlation of 0.92, temporal correlation of 0.76, interannual correlation of 0.69, and grid-cell level correlation of 0.6, between predicted and observed burned area. Our analysis reveals universal global patterns in fire-climate interactions, coupled with strong regional differences in fire-human relationships. Given the current socio-anthropogenic conditions, Equatorial Asia, southern Africa, and Australia show a strong sensitivity of fire extent to temperature whereas northern Africa shows a strong negative sensitivity. Overall, forests and shrublands, show a stronger sensitivity of burned area to temperature compared to savannas, potentially weakening their status as carbon sinks under future climate-change scenarios.

## 1 Introduction

Fires have been an integral part of the Earth system^1^ since the late Silurian c.420 ma^2^, while hominin-controlled fires have raged since the Middle Pleistocene c.700 ka^3^. Climate and human activity are thought to be the critical determinants of wildfire frequency, intensity and extent presently^1, 4, 5^. In turn, fires have not only shaped vegetation type at regional scales^6^ but can also cause abrupt shifts in vegetation state^7^. Although c.40% of the land area is fire-prone^6^, an average of c.3% of the land area has burned every year in recent decades, resulting in mean global carbon emissions of 2.2 PgC/yr which is c.25% of global anthropogenic C emissions^8, 9^. Although most natural wildfires are expected to be carbon neutral in the long run, the time required to sequester the burnt biomass may well run into several decades, especially in forest ecosystems. Repeated fires may further hinder sequestration, potentially resulting in positive net carbon emissions. Biomass-burning related greenhouse gas (GHG) as well as non-GHG emissions^10, 11^ and the changed post-burn albedo^12^ alter the atmospheric radiative balance, causing cascading effects on climate and vegetation^1, 13^. Wildfires also pose serious threats to human safety^14^.

Adequately characterizing the climate-human-vegetation-fire interactions is crucial to projecting the future of the Earth system, especially in the context of increasing human activity and ongoing climate change^15, 16^. Essentially, fires need sufficient fuel (biomass) in a flammable state (low moisture and high density), weather conditions suitable for enhancing fuel production, flammability, and fire spread, and a source of ignition (lightning, humans)^17^. Studies have suggested that, on the one hand, human influence is causing a decline in global burned area^18^ whereas, on the other hand, increasing global temperatures may lead to an increase in burned area in future^19–21^. Whereas the spatio-temporal effects of the physical drivers of fire are well characterized, those of anthropogenic drivers remain less understood.

At the global scale, biophysical process-based fire modules have been developed as components of dynamic global vegetation models (DGVMs)^22–25^, which have grown increasingly complex over time. However, despite their complexity and mechanistic appeal, their accuracy remains modest, with global spatial correlations between predicted and observed burned area in the range of 0.16−0.69 at a resolution of 1*° -* 2.5*°* ^26, 27^. Most models are also unable to predict the long-term decline in global burned area over the last two decades^18^. One reason for the lack of accuracy of global models may be, as we show in this study, that unlike fire-climate interactions, fire-human interactions may qualitatively differ between regions. In such a case, a better understanding of the human drivers of fire can come from an empirical framework that does not require any a priori assumptions regarding how humans influence fires.

Empirical approaches have been widely used to predict fire extent and to identify the drivers of fire^18, 28, 29^. However, non-linearities in fire-driver relationships pose a strong constraint on the accuracy of simple statistical models. A few studies that have accounted for non-linearities using more advanced statistical analyses^30, 31^ have been limited to specific regions or specific years. Empirical models usually also treat the spatial and temporal dimensions of fire extent separately, relying on either spatial or temporal correlations between burned area and socio-environmental variables to make inferences. Such a separation allows for analysis of drivers in space and time, but reduces the model utility for prediction of wildfires.

To address these twin issues of regionality and non-linearity, we use a machine-learning framework to understand the specific regional patterns of fire-climate-human interactions. We develop an artificial neural network model to predict burned area from socio-environmental drivers. Previously, studies have employed a similar approach^32–34^ to predict fire incidence probability. Here, we extend this approach with a simple yet novel neural-network architecture to directly predict burned area. To test how far a purely data-driven approach can go in predicting fire extent and to allow a comparison with other statistical models, we do not impose any mechanistic constraints on the model. Instead, the model learns fire-driver relationships exclusively from data. The machine-learning framework: a) can account for the high skew in the distribution of global burned area as well as the non-linearities in the fire-driver relationships, b) is sufficiently scalable to take advantage of large climate and socio-economic datasets which have become available, c) achieves high predictive accuracy with the least number of input variables, and d) can inform the parametrization of larger vegetation models.

## 2 Methods

### 2.1 Modelling approach and choice of drivers

A neural network is essentially a niche model, which not only delineates a volume in the space of drivers where fire occurs, but also predicts a value (in our case, burned area fraction) for each point in the driver space. Our choice of drivers broadly accounts for the classic factors that influence fire^17^: a) fuel biomass, b) its flammability, and c) ignition sources and fire management. A full list of variables used in our model can be found in SI-Table 2.

As a measure of fuel biomass, we use various cumulative measures of gross primary productivity (GPP). Fuel may comprise of litter, canopy, and grass. As a measure of litter biomass, we calculate total GPP over one year covering the previous growing season. In the northern hemisphere, this is essentially the previous calendar year, and in the southern hemisphere, the previous calendar year shifted by 6 months. As a measure of canopy and grass biomass, we use the cumulative GPP over one year up to the previous month. Not all the accumulated GPP will end up as fuel, especially due to differences in biomass allocation to roots, stem, and leaves. We do not explicitly model allocation, rather, indirectly account for allocation differences via vegetation type (as defined by the University of Maryland classification in the MODIS land-cover dataset; see SI-Table 2).

Fuel flammability depends on the intrinsic structural characteristics of the fuel and its moisture content. We use vegetation type to account for the differences in flammability and composition of fuel in different biomes. Moisture content of the fuel is accounted for by environmental variables, specifically, temperature, cloud cover, precipitation, and vapour pressure. Precipitation suppresses fire instantaneously, but over longer timescales, enhances fuel production and increases fire activity in the subsequent dry season. Previous studies have used various cumulative effects of precipitation to account for these long-term effects. Here, we only use the instantaneous values of precipitation, which has an effect of increasing fuel moisture and reduce flammability. The long-term effects are captured more directly via fuel proxies as described above.

Ignition sources are accounted for by human population density, cropland fraction, and lightning frequency. However, global monthly lightning data is not available for the full time period considered in this study. Therefore, we have used an aggregated dataset with global gridded mean monthly lightning climatology. Furthermore, an examination of pairwise relationships between drivers reveals that lightning frequency is strongly (but non-linearly) related to one or more other drivers, such as precipitation and cloud fraction (SI-Fig. 2). Therefore, these drivers further act as a proxy for lightning in the temporal dimension.

Fire management depends on fire prevention and suppression activities by individuals as well as institutionalized mechanisms. Human population density and cropland fraction, apart from being ignition sources, are also expected to have a role in fire suppression. Additionally, we consider road network density as a proxy of accessibility of the region for fire management. We train separate models at a subcontinental scale to broadly account for differences in institutional mechanisms, cultural factors, and habits. Ultimately, the spatial delineation of regions must be fine enough to capture such differences, but broad enough to generate sufficient data for training. As a reasonable classification, we use the regions as defined in the Global Fire Emissions Database (to ensure that sufficient training data is generated for the models, we have combined regions with low geographical area or fire incidence: TENA+CEAM = TCAM, NHSA+SHSA = SA, EURO+MIDE = EUME).

SI-Table 2 lists all variables considered, along with the datasets used and any pre-computations performed on raw data.

### 2.2 Choice of spatial resolution

To unify the temporal and spatial dimensions, we need a spatial resolution at which the negative effect of past fires on current burned area is low. Therefore, we choose a coarse spatial resolution of 1*° ×* 1*°*. At this scale, as long as burned fractions are low, new fires can still occur in other parts of the gridcell which were not previously exposed to fire, diminishing the overall effect of fire history. Fortunately, this also works for grid-cells with high burned fractions, because such cells are typically located in African savannas, which replenish fuel every year. To verify this reasoning, we built a null model that predicts present burned area only from annual fire history, and found a strong positive correlation (high predictability) between present fire and fire history. This confirms that at this spatial scale, fire history merely reflects the combined effects of other fire drivers without the negative temporal effect.

Vegetation type can change at a much finer spatial scale, especially in areas fragmented by croplands. To account for these fine-scale variations, we calculate the fractional area under each vegetation type in each grid-cell from a high-resolution (MODIS Land Cover) dataset, rather than using a single dominant type at the cell level. The fraction of each vegetation type is then used as an input to the neural network.

All data was coarse-grained to 1*° ×* 1*°* resolution (average of all data-points falling within each 1*° ×* 1*°* grid-cell). For model training and analysis, we only consider gridcells with at least 30% natural vegetation.

### 2.3 The neural-network model

We feed the input variables (SI-Table 2) into a dense neural network with a single hidden layer consisting of 12 neurons and ELU activation. The output layer consists of 25 neurons with softmax activation. To account for the high skew in the burned area distribution, we divide the burned area range [0, 1] into 25 intervals (classes). The first interval is [0, 10^−6^) and the remaining 24 intervals divide the range [10^−6^, 1] equally on a log scale. Each output neuron predicts the probability *p*_*i*_ of burned-area being in class *i*, from which we calculate actual burned area as *BA* = Σ_*i*_ *p*_*i*_*B*_*i*_, where *B*_*i*_ is the geometric mean of the bounds of class *i*.

We train the model using the GFED4.1s burned area dataset^9^, which specifically accounts for small fires neglected in earlier datasets. We divide our data into training, evaluation and test datasets, and train the network by minimizing cross-entropy on the training dataset. We halt training when prediction accuracy converges on the validation dataset. We evaluate the performance of different alternative models on all data, which includes the test dataset. We use monthly data from 14 years between 2002-2015 for our analysis. Of these, all data in years 2005-2007 is designated as the test-data. From the remaining data (all grid-cells for all months except 2005-2007), a random sample of 70% of the data points is used for training, and the remaining 30% data points are used for validation. To minimize overfitting, we keep the number of neurons in the hidden layer to a minimum, such that no substantial accuracy is gained from further increasing it. The code to format data and run the Neural-Network model is publicly available at https://github.com/jaideep777/neuralfire.

### 2.4 Measuring model performance

To rank models by performance, we calculate five performance metrics for each model - temporal and interannual correlations between spatially aggregated monthly and yearly timeseries of burned area (*r*_*T*_ and *r*_*IA*_), correlation between predicted and observed yearly anomalies (*r*_*An*_), spatial correlation between mean yearly burned area (*r*_*S*_), and fractional deviation of predicted total yearly burned area from that observed (*r*_*BA*_ = 1 − abs(1 −*<u>BA</u>*_predicted_*/BA*_observed_)). We then combine these metrics with weights (*w*_*i*_) into an aggregate performance score 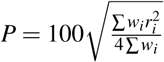, which ranges between 0 − 100, higher the better. These metrics are not used in NN training, but only to rank trained models (i.e., the models are optimized for cell level, and not aggregate, performance). We aim to identify models that have good inter-annual predictability. However, since the spatial extent of data is much greater than its temporal extent, if all weights were equal, models that perform well spatially would receive a higher score even if they delivered poor interannual predictability. Therefore, to privilege models with better interannual predictability, we use *w*_*IA*_ = 4 and all others weights *w*_*i*_ = 1. We report the correlation between predicted and observed BA in individual grid-cells (*r*_*I*_, SI-Fig. 1), but do not account for it in evaluating the model performance. This is because we found *r*_*I*_ to be a poor indicator of model performance: even simple linear models perform very well on this metric even though they get the spatio-temporal fire patterns as well as total burned area completely wrong.

For each region, we begin with training a model that uses all socio-environmental variables as predictors. Then, we drop one or more variables, trying out different combinations of drivers and measuring the model performance *P*, until we arrive at the best performing model. We then further drop variables to arrive at a ‘minimal model’, i.e., a model that uses the least number of variables without a substantial performance loss compared to the best model (we use the criterion, *P*_*best*_ *-P*_*minimal*_ *≤* 3). For global analysis, we mosaic predictions from the minimal regional models for each timestep.

## 3 Results

### 3.1 Regional differences in the observed fire niche

To lay the ground for visualizing our model results and justifying our modelling framework, we first plot the fire niche from observed (GFED4.1s) data. The fire niche can be thought of as an n-dimensional function, where each dimension is a socio-environmental variable and the function value corresponds to burned area. This fire niche can be further understood by looking at the density distribution of the drivers in the same n-dimensional space, where the density corresponds to the number of times that driver combination occurs. We show how seemingly similar biomes can have very different fire regimes due to differences in human activity.

First, we find that weather imposes universal constraints on fires: fires are limited when temperatures are very low or very high, occurring largely at temperatures between 15 − 30*°*C (Fig. 1). Similarly, high (instantaneous) precipitation suppresses fires, with most fires occurring when precipitation is below 5 mm/month (Fig. 1). The intermediate productivity hypothesis^35, 36^ can also be clearly observed, as fires occur for intermediate values of cumulative GPP across all regions (Fig. 2).

**Figure 1.**
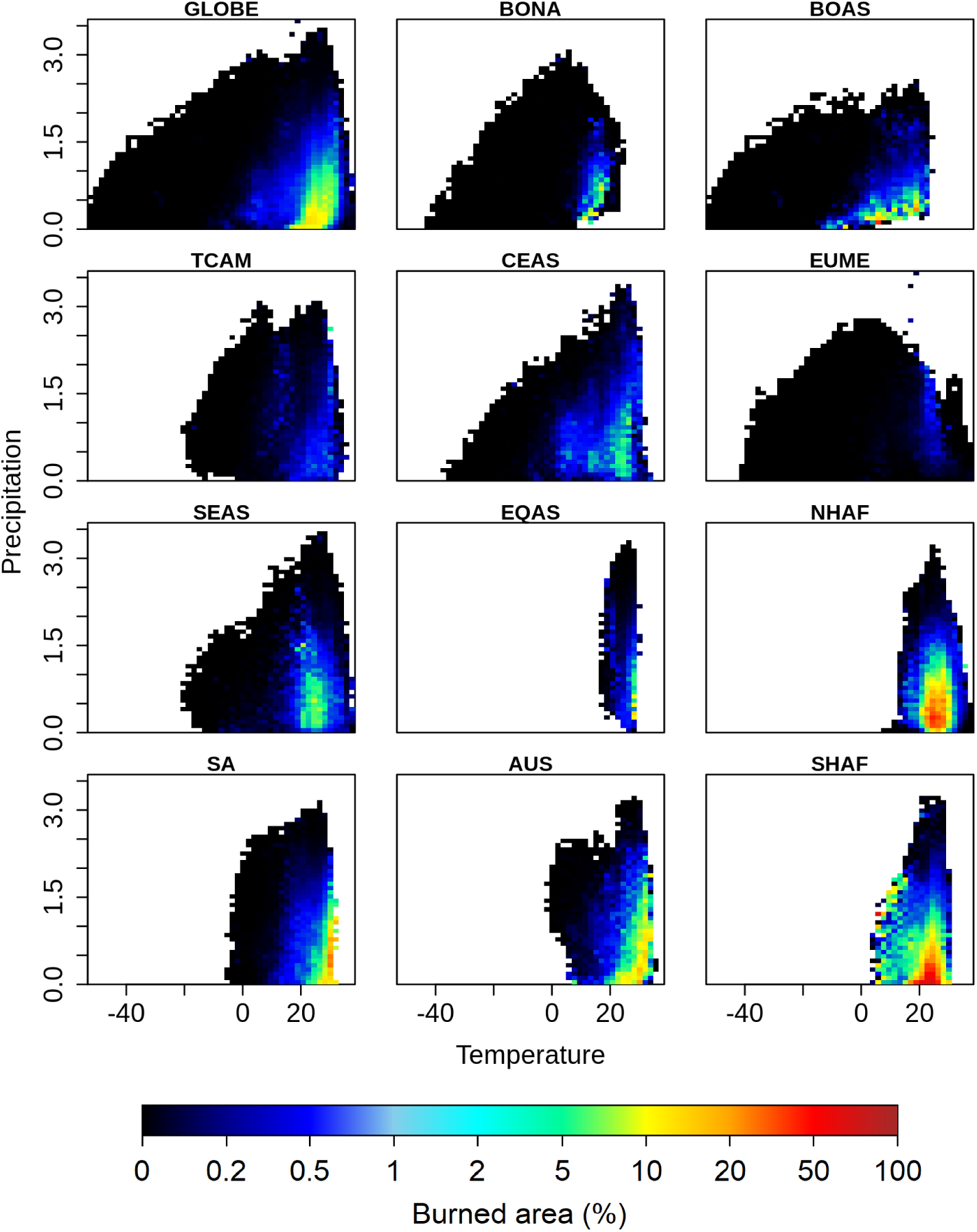
Visualization of the fire niche along the temperature-precipitation axes. A) Each coloured (non-white) point represents a realized value of the driver pair, with the colour indicating mean burned area fraction observed for that driver pair. Here the precipitation axis is log transformed with the function *y* = log(1 + *x*). Fires are largely confined between 15 − 30*°*C temperature and < 5 mm/month precipitation (about 1.5 units on the log transformed scale shown here). Notable outliers can be seen in southern hemisphere Africa, where fires are observed at lower temperature and higher precipitation.

**Figure 2.**
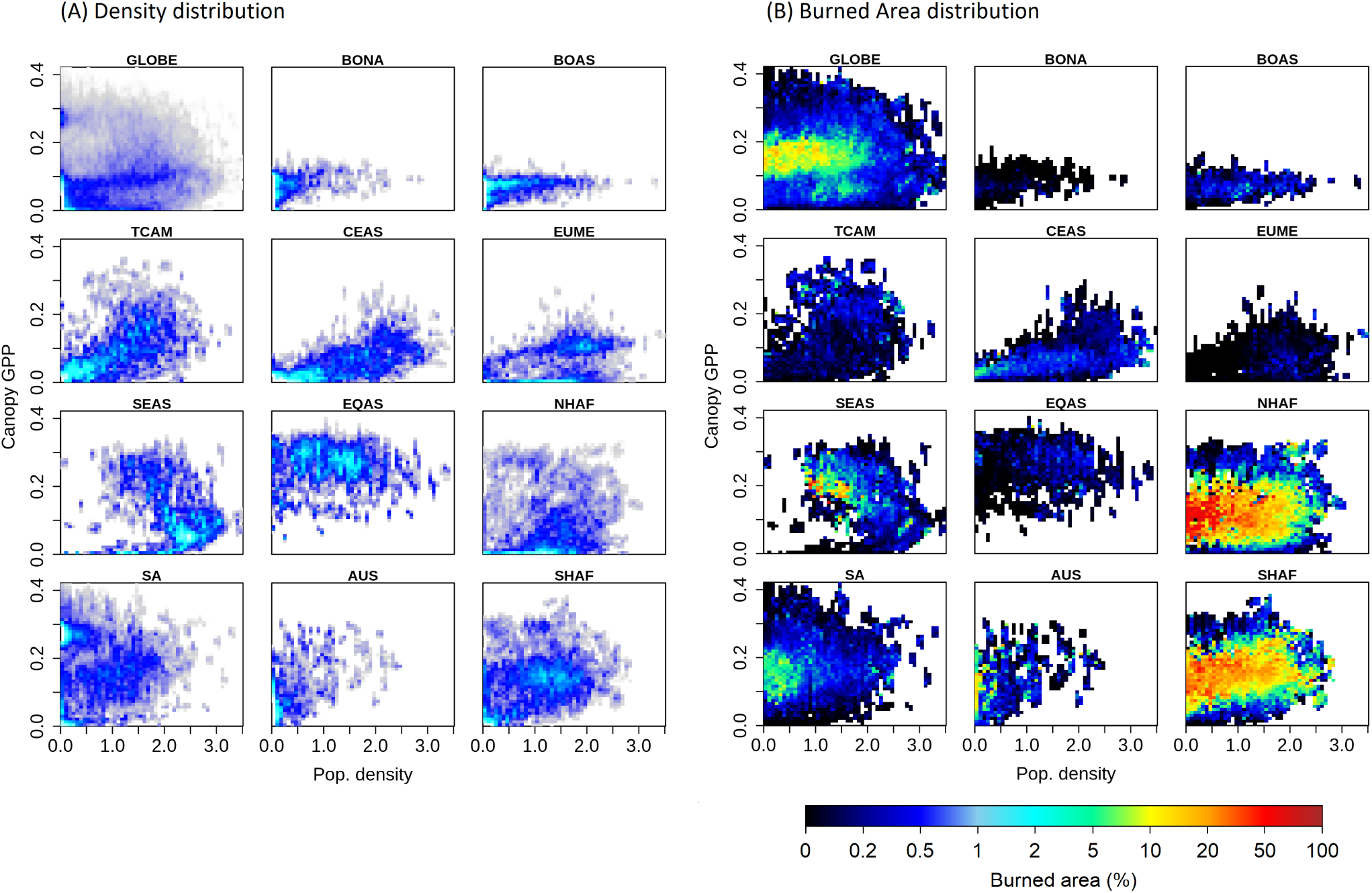
Regional differences in fire regimes can be seen along the GPP-population density axis. The frequency of occurrence of different GPP-population density driver pairs (A), and the mean burned area observed for each pair (B). Population density is log-transformed with the function (*y* = *log*(1 + *x*)). In general, fires occur at intermediate values of GPP and decrease with population density. However, the responses of burned area to population density are starkly different in different regions: In South America, burned areas are already low at low population densities (likely due to lower temperatures experienced there), and decrease sharply to almost zero once population density crosses about 3 persons/km^2^. By contrast, fires persist till very high population densities and decline only gradually with increasing population density in Africa. In Australia, burned areas are high at near-zero population densities, but decline sharply even for small population densities.

However, strong regional differences can be observed when we look along anthropogenic dimensions, where similar environmental conditions lead to very different fire regimes. For example, the responses of burned area to population density differ strongly between regions - burned area declines sharply with population density in Australia and South America, declines gradually and persists until much higher population densities in Africa, and even increases with population density in Boreal and Equatorial Asia. Indeed, the most surprising difference is between northern and southern Africa. Despite having similar environmental conditions, similar biomes and phylogenetically similar vegetation, the fire niches in the two regions are substantially different: in northern Africa, we find a high fire extent in areas with low GPP (marked with a white triangle in Fig. 2), but not so in southern Africa; similarly, we find high burned area in southern Africa in regions with high population density and high GPP, but not so in northern Africa (marked with white ellipse in Fig. 2). This makes a solid case for a separate treatment of different subcontinents in fire modelling.

### 3.2 Model performance

We now compare the predictions of regionally trained neural-network models with the observed data. At the global scale, predictions from our model (mosaicked regional models) closely match observations (Fig. 3; see SI-Table 1 for the complete set of models and their regional performance statistics). Our model accurately captures the spatial, seasonal, and interannual variability in burned area, with correlations between predicted and observed data as follows: spatial correlation using temporally averaged burned area - *r*_*S*_ = 0.92, temporal correlation using global monthly burned area - *r*_*T*_ = 0.76, interannual correlation using global yearly burned area - *r*_*IA*_ = 0.69, and individual correlation (between burned area of individual gridcells across time and space) - *r*_*I*_ = 0.6 (SI-Fig. 1). We also evaluated the performance of our model among different vegetation types, in which *r*_*I*_ varies between 0.31 − 0.78. The model performs best in savannas and broadleaved evergreen forests, and worst in closed shrublands and needleleaved forests (SI-Fig. 1). Further, our model predicts a total burned area of 460.41 Mha, against an observed value of 445.26 Mha.

**Figure 3.**
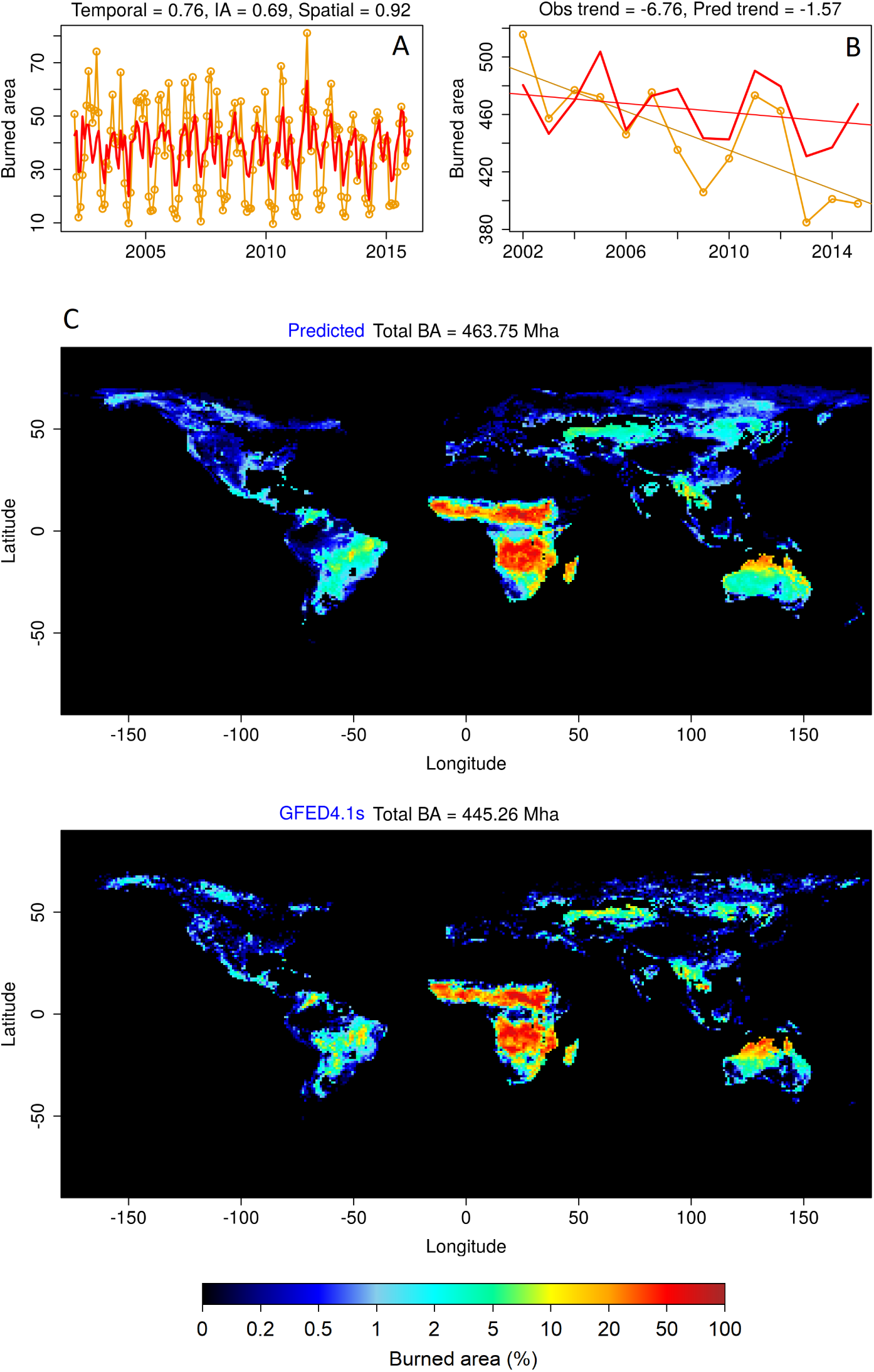
Spatio-temporal performance of our model at the global scale. Temporal monthly (A), Annual (B), and Spatial (C) burned area predicted by our model (solid red lines) compared with the GFED4.1s burned area data (orange lines and circles). Our model accurately predicts the spatial distribution of fires across the globe, with a spatial correlation of 0.92. It captures the yearly anomalies in global burned area reasonably well (with temporal correlation of 0.76 and interannual correlation of 0.69), and predicts a long term decline of 1.57 Mha/yr^2^ during the years 2002-2015, compared to an observed decline of 6.76 Mha/yr^2^. predict a negative trend closer to the actual, but all three substantially underestimate mean global burned area (< 350 Mha/yr)^18^.

Models trained for different regions vary in their performance. The best predictability (performance score *≥* 95) is achieved for fires in Equatorial Asia, Australia, and South America, whereas those in Boreal regions, Europe, and the Middle East are the least predictable (score between 74 − 85) (Table 1). Model predictions of interannual fire patterns are best in regions with frequent fires (*r*_*IA*_ range between 0.68 − 0.89). *r*_*IA*_ is lowest in regions with rare fires, especially in Boreal regions (0.56 − 0.59). Our models suffer from a slight positive bias in regions which contain very high burned fractions, such as in interior Australia and equatorial southern Africa. In such regions, our model predicts a burned area of about 2.5 % in cells with extremely low burned area (< 1%).

**Table 1.**
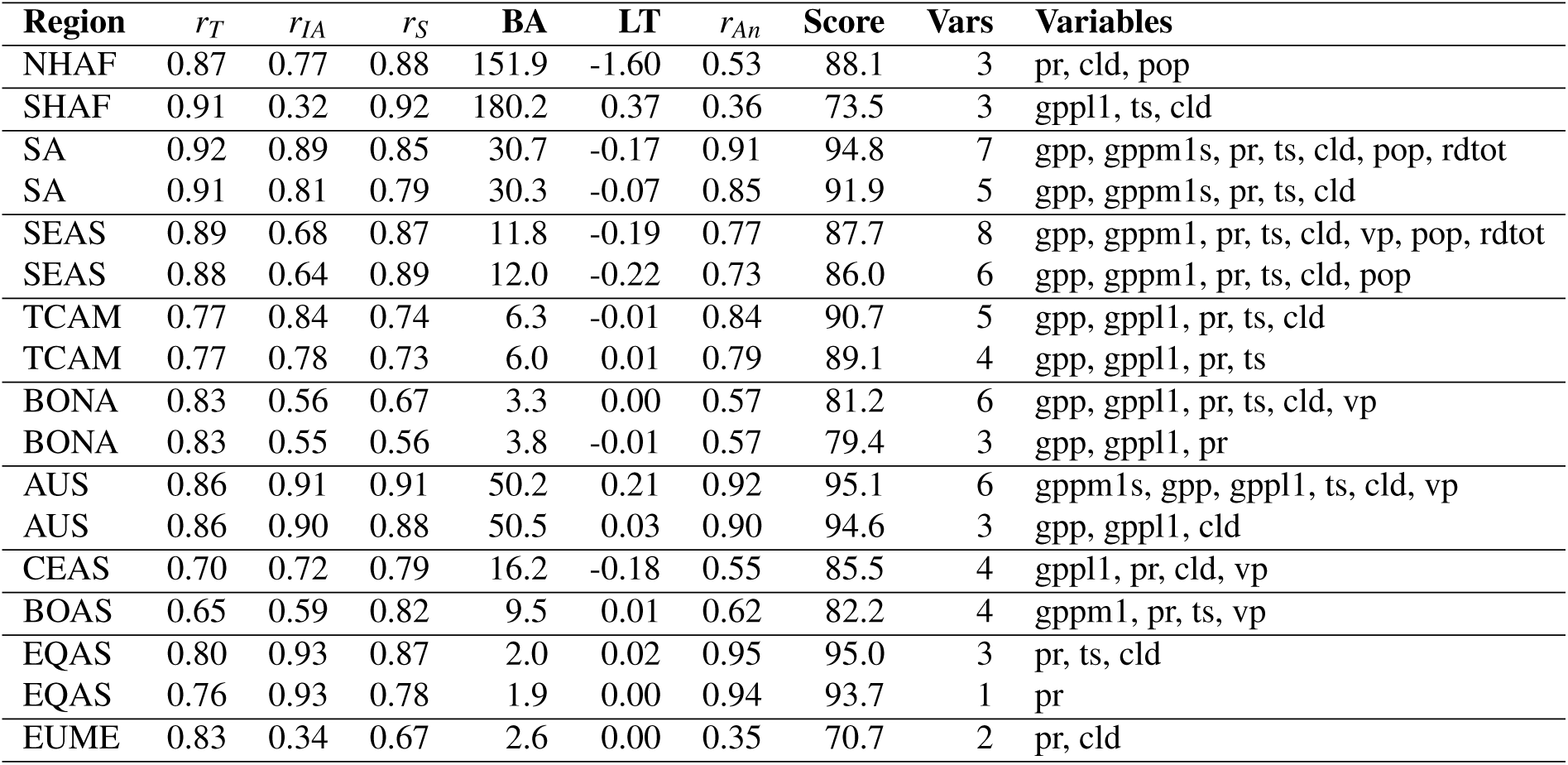
Regional predictors of fire. Performance of the best and minimal models for each region with respect to each of the five performance measures described in Methods, along with the aggregate performance score. In some regions, the best model is the same as the minimal model. Also mentioned are the variables that form the inputs of the models. BA is Burned Area, and LT is long-term trend in spatially aggregated yearly timeseries. Variables are as follows: gppl1 - cumulative GPP, gppm1 - growing season GPP (northern hemisphere), gppm1s - growing season GPP (southern hemisphere), pr - precipitation, ts - temperature, cld - cloud cover, vp - vapour pressure, rdtot - total road network density, pop - population density. All models include vegetation type fractions, including cropland fraction. The model for NHAF uses yearly vegetation fractions, whereas rest of the models use a single snapshot.

Figure 4 compares the predicted and observed interannual fire extent and long term trends for each region. Although 15 years of data are too short to train our model to capture long-term trends, our model captures *∼* 23% (1.57 Mha/yr^2^ of 6.76 Mha/yr^2^) of the observed global decline in burned area (derived from GFED4.1s data). More than 60% (4.13 Mha/yr^2^) of the observed global decline is contributed by northern Africa, out of which our model captures 39% (1.60 Mha/yr^2^) with dynamic vegetation fractions (SI-Table 1), and 36% (1.51 Mha/yr^2^) otherwise (Fig. 4D). By contrast, only three of seven process-based models from the FireMIP project running at a sub-yearly temporal resolution (CLM fire module, MC-Fire, and JULES-INFERNO)

**Figure 4.**
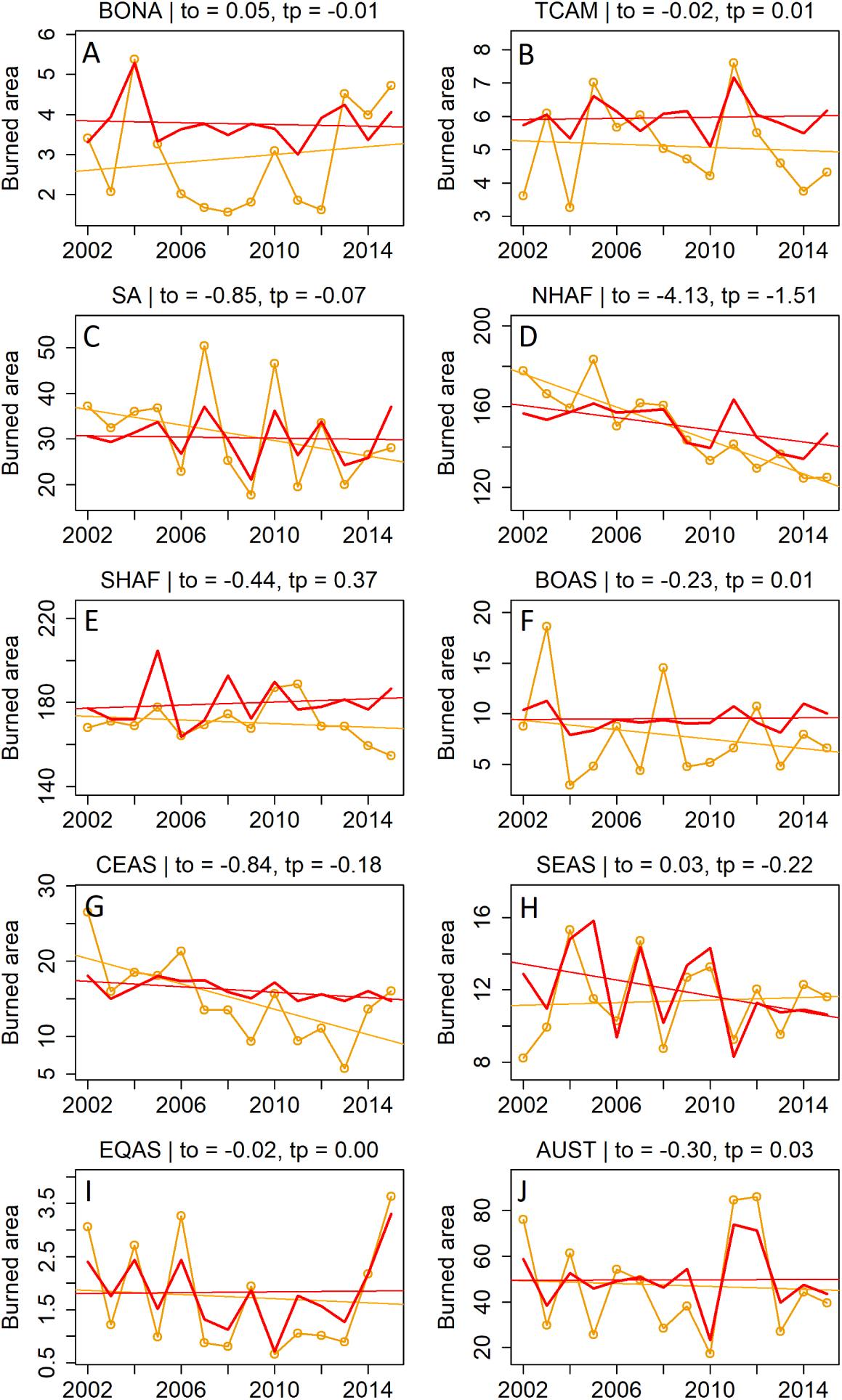
Predicted and observed interannual fire extent. Total annual burned area observed in different geographic regions (orange lines and circles) along with that predicted by the minimal model for each region (solid red line). Regression lines indicate the long-term trend in burned area, with the trends in observed (*t*_*o*_) and predicted (*t*_*p*_) burned areas mentioned above each panel. Interannual variability in burned area is well captured in our model, especially in Equatorial Asia, Australia, Southeast Asia, and South America. Long-term decline is highest in northern Africa, with our model predicting 36% of the observed decline. Fires in southern Africa drop sharply after 2013.

### 3.3 Sufficient regional predictors of fire

For each region, we obtain the sufficient regional predictors of fire from the inputs of the regional minimal models. Regional sufficiency of socio-environmental factors does not necessarily imply that other factors are not of mechanistic importance in fire ignition and spread. A factor that clearly favours fire may drop out of a regional model if, a) it is not sufficiently variable in the subcontinental region (e.g., fuel load is always high in Equatorial Asia, and population density is always very low in Australia), or b) if it is correlated with another factor that influences fire (e.g. in Australia, either of temperature and cloud cover is sufficient to predict fire because both are correlated, but temperature drops out of the minimal model for Australia). On the contrary, factors that are not necessarily limiting (such as fuel load in Boreal regions) may still be significant predictors due to variability within the region. The sufficient predictors identified here should thus be interpreted as those which have the highest predictive value at the subcontinental scale, given the sum total effects of all socio-environmental drivers.

Within each region, climate and fuel load explain the spatio-temporal patterns of fire extent for all regions except northern Africa and Southeast Asia, where human population density is additionally a significant predictor (Table 1). Fuel load turned out to be a significant predictor in all Boreal regions in our study, whereas it was not considered to be significant in previous studies (SI-Table 4). Vegetation type fractions explain most of the spatial variability in fires across all regions, whereas climate and fuel load were the most important predictors of seasonal and interannual variability. Among the anthropogenic factors considered, population density had a negative effect on fire, with a monotonic decline in burned area with increasing population density, but within-region variability in population density was important only in northern Africa and Southeast Asia.

To test the effect of cropland fraction, we excluded two vegetation type fractions from model training (fraction of area under croplands as well as the fraction of non-vegetated area from the minimal model for each region), and compared the resultant models with the original minimal models. For this analysis, it is not enough to exclude only cropland fraction: as vegetation-type fractions add up to one, excluding any one fraction still provides the neural network with all land cover information. Cropland fraction was a significant predictor in Southeast Asia and Boreal North America (i.e. predictability reduced when cropland fraction was excluded in these regions). In Boreal Asia and Central Asia, exclusion of cropland fraction improved predictability, implying that cropland fraction is neither a consistent driver nor a consistent deterrent of fires in these regions.

Road network density and lightning climatology showed no substantial explanatory power within regions, and dropped out of all regional minimal models (for the effects of lightning, compare version 8 models in SI-Table 1). However, data on both these variables were not available for multiple years. Therefore we do not rule out their effect on fires based on this study. Specifically, including monthly lightning data may improve predictions in Boreal regions, as these regions are known to be frequently ignited by lightning.

Droughts associated with El Niño events have been shown to strongly influence fires across the tropics, especially South America and Equatorial Asia^37^. Higher fires associated with El Niño events are observed in South America in the years 2007, 2010, and 2015 (Fig. 4C)^38^, and in Equatorial Asia in 2002, 2004, 2006, 2009, and 2015 (Fig. 4I)^39^. Our model correctly predicts high burned area in these regions and years. Furthermore, the extreme fire events observed in Australia in 2011 and 2012 appear to be caused by negative values of the Interdecadal Pacific Oscillation (IPO) coupled with El Niño (negative values of the Southern Oscillation Index)^40, 41^. Our model also predicted high burned areas in Australia during these years (Fig. 4J).

Previous studies have attributed the long-term decline in fire in northern Africa to cropland expansion^42^. However, we find that this decline is instead explained most strongly (39%) by increasing population density. We found no performance drop after excluding cropland fraction from the model (compare version 6 models in SI-Table 1), implying a low predictive value of cropland expansion. The residual long-term decline in northern Africa does not appear to be driven by changes in climate or vegetation either - we did not achieve better predictive ability even after including dynamic (yearly) vegetation fractions in our training (SI-Table 1), even though trends in certain vegetation type fractions are weakly correlated with trends in burned area (SI-Figs. 4 and 5).

Our models were also able to broadly distinguish fuel characteristics in different regions. In Africa, Australia, and Central Asia, the cumulative GPP up to the previous month (a proxy for grass biomass) featured in the minimal models, whereas in Southeast Asia the previous calendar year’s GPP (a proxy for litter biomass) did. In South America and Australia, both litter and canopy biomass were equally good predictors of burned area, implying that both might constitute the fuel in those savannas.

### 3.4 Climate Sensitivity

Given the current socio-environmental conditions, how will different regions respond in terms of wildfire vulnerability to increasing global temperatures? Towards answering this question, we drive the best regional model for each region with the same input data from the time period 2002-2015, but with temperature uniformly incremented by a small amount (Δ*T*), while keeping other variables at their original values. This small change in temperature is assumed to have no effect on the vegetation type distribution. In case temperature has dropped out of the best regional model, we choose the next best model that includes it (SI-Table 3 lists the models used in such cases). We then measure the sensitivity of burned area to temperature as the change in burned area fraction per unit change in temperature (*S* = Δ*BA/*Δ*T*), and as percent change per unit temperature (*S*_%_ = Δ*BA/BA/*Δ*T*).

Globally, forest-dominated areas show the highest sensitivity 10.46% − 18.75%*/°C* of burned area to temperature (Table 2). Grasslands and croplands show moderate sensitivity (4.91%*/°C* and 8.65%*/°C* respectively), whereas savannas show a negative slight sensitivity (− 0.57%*/°C*). In absolute terms, the most sensitive areas are concentrated in Equatorial Asia, southern Africa, and northern Australia. This is a result of high sensitivity during the months of August-November (Winter-Spring), when fires are currently temperature limited (Fig. 5B). Northern African savannas show a strong negative sensitivity to temperature (Fig. 5A), with the effect being strongest in the months of February-April (summer) and weakest in December-January (winter), with some areas even showing a positive sensitivity in winter. Therefore, this decrease is likely due to a reduction in biomass density associated with an increase in aridity (vegetation type is held constant). Southeastern Australia and eastern Himalayan regions have relatively less fires, but are highly sensitive to temperature changes in terms of percent change in burned area (SI-Fig. 6). SI-Fig 3 (animated gif) shows the global sensitivity for each month.

**Table 2.**
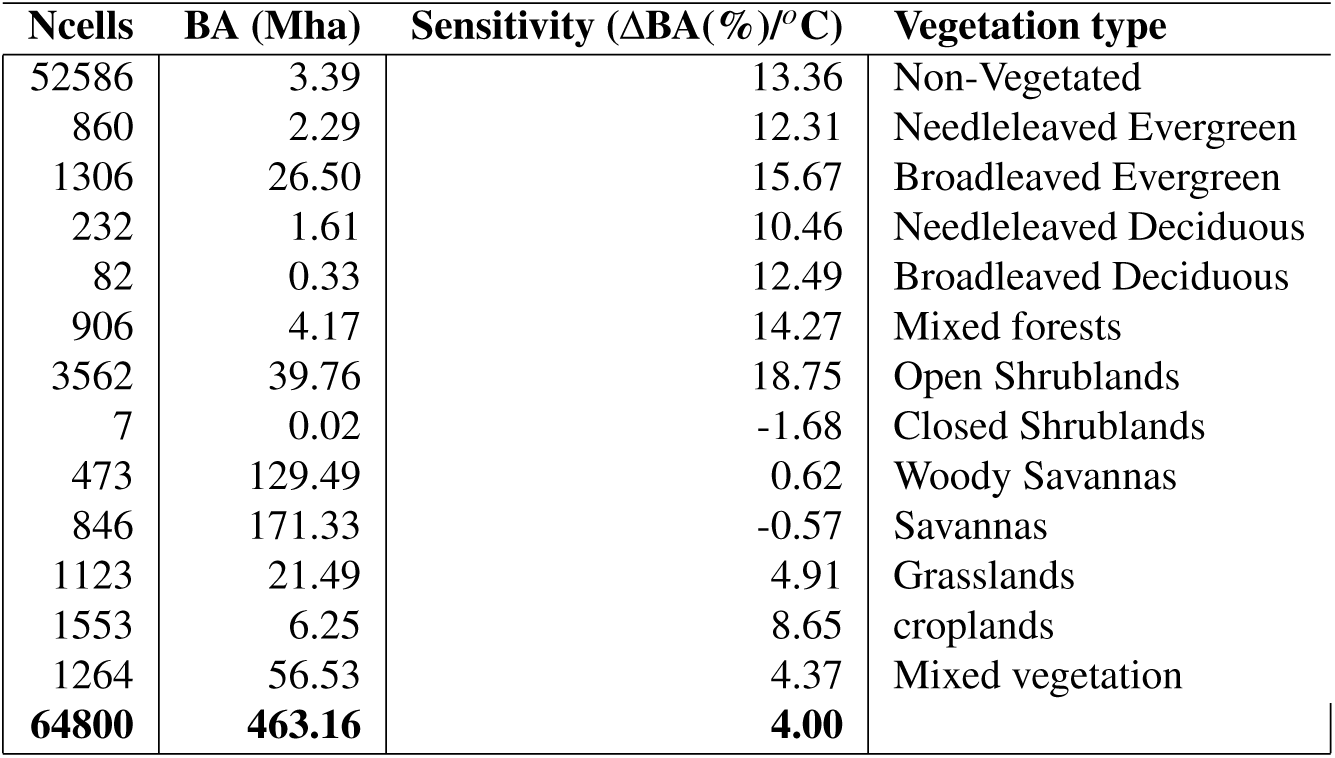
Sensitivity of different vegetation types to increase in temperature. Compared to a decrease in savannas of 0.57%, there is a disproportionately positive sensitivity of burned area to temperature in forests and open shrublands. Numbers in bold indicate global totals.

| Ncells | BA (Mha) | Sensitivity (ΔBA(%)/$^{\circ}$C) | Vegetation type |
| --- | --- | --- | --- |
| 52586 | 3.39 | 13.36 | Non-Vegetated |
| 860 | 2.29 | 12.31 | Needleleaved Evergreen |
| 1306 | 26.50 | 15.67 | Broadleaved Evergreen |
| 232 | 1.61 | 10.46 | Needleleaved Deciduous |
| 82 | 0.33 | 12.49 | Broadleaved Deciduous |
| 906 | 4.17 | 14.27 | Mixed forests |
| 3562 | 39.76 | 18.75 | Open Shrublands |
| 7 | 0.02 | -1.68 | Closed Shrublands |
| 473 | 129.49 | 0.62 | Woody Savannas |
| 846 | 171.33 | -0.57 | Savannas |
| 1123 | 21.49 | 4.91 | Grasslands |
| 1553 | 6.25 | 8.65 | croplands |
| 1264 | 56.53 | 4.37 | Mixed vegetation |
| **64800** | **463.16** | **4.00** |  |

**Figure 5.**
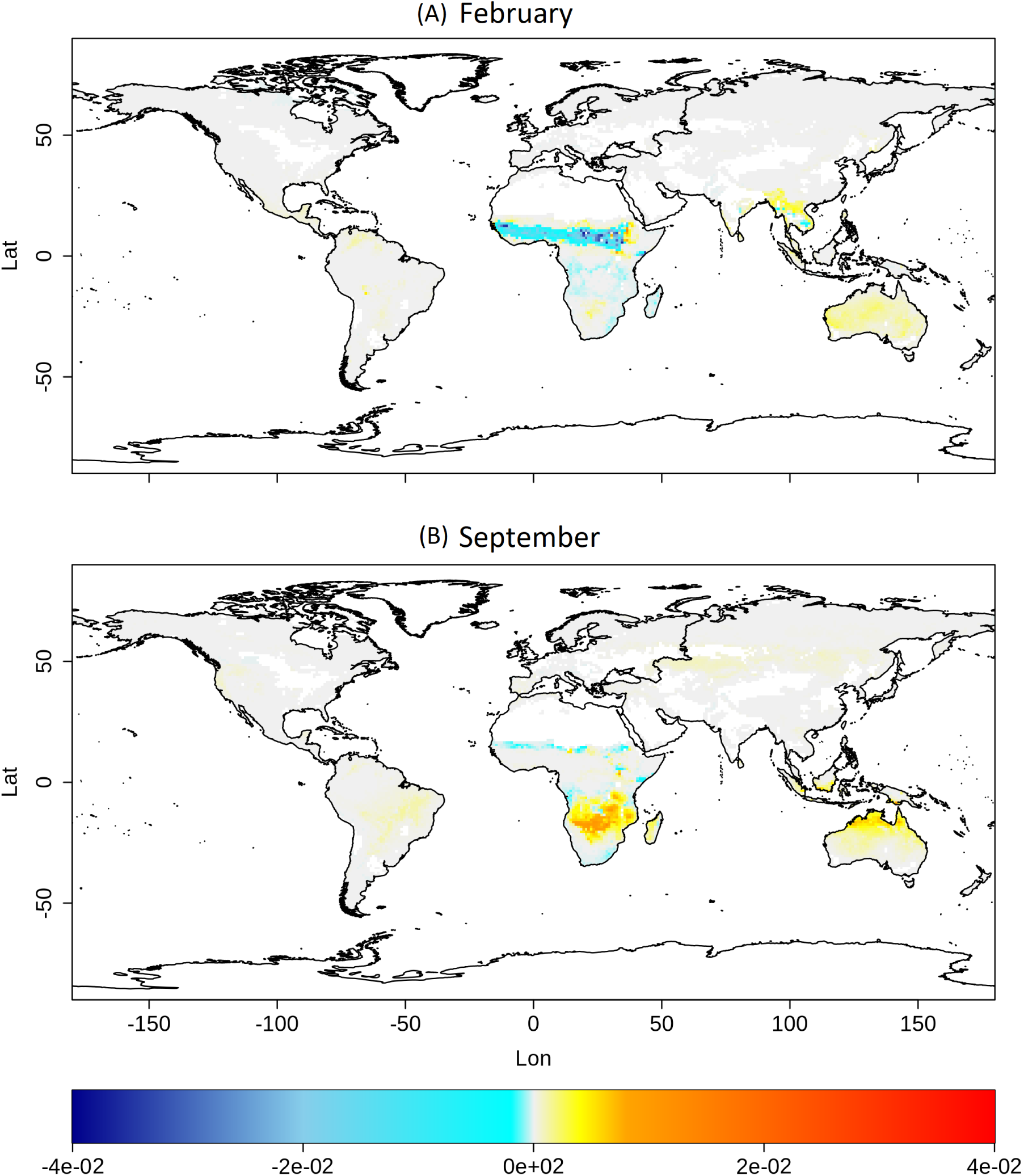
Sensitivity of burned area to temperature. Change in burned area fraction per *°*C rise in temperature in February (A) and September (B). While incrementing temperature, all other variables were held equal to their original values.

## 4 Discussion

Our machine-learning model rivals more complex process-based models, and delivers higher accuracy with fewer input variables. We found distinct regional differences in fire drivers across regions, but within regions, between 1-5 drivers are sufficient to accurately predict burned area. We found that whereas climatic constraints on fires were universal, differences in anthropogenic niches may drive regional differences in fire activity. We predicted differential effects of increasing temperature in different regions, with forests being disproportionately sensitive to temperature changes compared to savannas, although we have not accounted for changes in co-varying drivers in this analysis. Our work suggests that an improvement in predictive accuracy of fire models can result from better parameterization of models with fewer drivers, rather than expanding already complex models with more processes and parameters.

Modelling approaches based in machine learning often face the criticism that they do not provide any understanding of the underlying mechanisms and processes. However, as we demonstrate in this work, it is now possible (due to advantages in computational power) to scale up neural-network models and run them iteratively to perform an analysis of the minimal predictors of fire. Such an analysis provides vital information on the relative importance of different drivers in different environmental conditions. Furthermore, it is possible to look into the functional relationships between fire and the most important drivers learned by the model, to make inferences about the underlying mechanisms as well as to parametrize process-based models. Although neural-network approaches have been previously used for fire incidence prediction^32–34,43,44^, our model uses a novel architecture (a multiclass classifier with fine-grained burned area classes on a log scale) to predict continuous burned area at continental and global scales.

Our model does predict cell-level extreme burned fractions with good accuracy (lesser spread towards higher burned fractions in SI-Fig. 1), but fails to distinguish fire extremes at an annual regional scale. For example, the extreme fires in Boreal Asia in 2003, 2007, and 2010 are not captured, whereas high burned area is predicted in Boreal America even in years with low fire activity (2006-2012). Fires during peak years in Equatorial Asia and South America are also slightly underestimated. This might be due to the rarity of extreme events, such that most of the training data consists of non-extreme fires, and an imbalance in the spatial and temporal extents of the training data. Due to the flexibility of the neural network approach, it is possible to assess the drivers of extreme fire events by training a model with a subset of data containing a greater proportion of extreme burned fractions. A related problem is that long-term annual decline is not fully captured by the models. To mitigate these issues, future work could use data resampling to equalize the spatial and temporal extent of training data.

Studies differ on the predicted drivers of fire for the same regions. For example, the drivers of fire in northern Africa are predicted to be precipitation, population density, and cropland fraction^42^, or population density, temperature, and wet days^31^.

We find precipitation and population density to be important drivers, but no effect of cropland fraction. In southern Africa, predicted drivers are fuel and climate^42^, or wet days and cropland fraction^31^, or tree-cover, rainfall, dry season, and grazing^30^. We, too, find fuel and climate to be the key drivers. While other studies predict fuel and climate to be the drivers in Equatorial Asia^29^, we found that precipitation alone explained the variability in fires in this region. Similarly, Abatzoglow et al. ^29^ find aridity alone as a driver in Southeast Asia, whereas we find climate, fuel, as well as population density to be important drivers. In Boreal areas, we find an important effect of fuel load, not predicted by previous studies. SI-Table 4 gives a comparison of predicted drivers in all regions.

Researchers agree that under future climate scenarios, some areas of the globe will see increased fire activity while others see a decline^4^. The contrast in fire sensitivity to increasing temperature between northern and southern Africa may seem surprising due to similarities in weather and vegetation. However, global climate models agree that fires in northern Africa may decline by end of the century, whereas model agreement for southern Africa is low, and on average models predict no change^45^. As we have argued, this could be the effect of differences in anthropogenic niche of fires in the two regions, with rather anomalous niches in southern Africa. To further confirm the predicted decline in fires in northern Africa, we ran the sensitivity analysis with models including and excluding human population density, and obtained similar results. Models also show scarce agreement on the change in fires in southeastern Australia, where our model predicts a high temperature sensitivity (in terms of percentage change). Qualitatively, our prediction may be corroborated by the recent occurrences of large-scale fires in this region during December 2019 to January 2020. One region where our model disagrees with the consensus of global models is northern Australia, where we predict an increase in winter-spring fires with temperature whereas global models predict and agree on a decrease. However, this discrepancy could be explained by accounting for precipitation, which is expected to increase in this region, but is assumed constant in our analysis. Furthermore, a small predicted increase in fires in already-arid interior Australia is also surprising, but is consistent with the consensus of global models, and could be an artefact of data-limitations as we argue below. A quantitative analysis of changes in burned area using future projected climatic drivers would provide more accurate projections of fire activity under future climate change scenarios. Our neural-network model can be directly integrated into vegetation models for such analyses.

We caution readers in interpreting the sensitivity values in arid areas in interior Australia and parts of South America. Although we expect fires to reduce at extremely high temperatures due to declines in vegetation cover, in Australia and South America, data-points which show a reduction in burned area at higher temperatures are limited (Fig. 1). Therefore, the neural network does not have the opportunity to learn this declining trend. This is in contrast with other regions which do show a decline in fires for extremely high temperatures. Therefore, the models might overestimate sensitivity to temperature in very arid areas within these regions. This problem could be overcome as more data becomes available.

Fuel consumed in fires and subsequent emissions vary by region^9, 46^. In savanna-dominated Africa, fuel consumption per unit area burned is low, and fires are carbon neutral as most of the grass biomass is regenerated the next year. By contrast, in tropical and Boreal forests, fuel consumption is high, not only from standing vegetation, but also from soil carbon, and recovery of lost biomass may take decades. On the one hand, burned area has declined due to human influence in African savannas, leading to an overall decline in global burned area. On the other hand, fires in Boreal and tropical forests are driven by climate, potentially putting these regions at a heightened fire risk due to future climate change. An increase in burned area in these forests may further increase fire related emissions, weakening their status as carbon sinks, and creating a cascading effect on the global climate system.

Our model does not attempt to capture the complex feedbacks from fire into the climate systems^47^ which, in any case, is a limitation even in advanced DGVMs^16^. Neural network models have the potential to be coupled with DGVMs and socio-economic drivers, and may provide a simpler class of models to predict future fire regimes, assess impacts such as GHG and non-GHG emissions, distribution of vegetation types, and risks to society, both at regional and global scales.

## Acknowledgements (not compulsory)

RS was a JC Bose National Fellow during the tenure of this study. Partial funding for the model development came from the Ministry of Environment, Forests and Climate Change, Government of India, as part of the country’s Third National Communication to the UNFCCC.

## Author contributions statement

JJ and RS designed the study, JJ performed the analysis, JJ and RS interpreted the results, JJ and RS wrote the manuscript, RS acquired funding. All authors reviewed the manuscript.

## Supplementary Information

**SI-Table 1.**
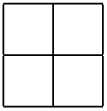
Please see SI-Table 1 excel sheet in Supporting Information.

**SI-Table 2.** Gridded datasets used and their soures

**SI-Table 1.** Please see SI-Table 1 excel sheet in Supporting Information.
|  | Variable | Data source | Spa. Res | Time Res | Details | Citation |
| --- | --- | --- | --- | --- | --- | --- |
|  | **Dynamic variables** |  |  |  |  |  |
| 1 | Temperature | CRU TS4.01 | 0.5° | monthly | Mean monthly temperature | 48 |
| 2 | Vapour pressure | CRU TS4.01 | 0.5° | monthly | Mean monthly vapour pressure | 48 |
| 3 | Cloud cover | MODAL2 / MOD06 | 0.05° | monthly | Mean monthly cloud fraction, coarse-grained |  |
| 4 | Precipitation | GPCP2.3 | 2.5° | monthly | Monthly total precipitation, transformed as $\log(1+x)$ | 49 |
| 5 | Current GPP | MOD17A1 | 0.05° | monthly | Mean monthly GPP | 50 |
| 6 | Growing season GPP | MOD17A1 | 0.05° | monthly | For northern Hemisphere: Sum of monthly GPP over 12 months of the previous calendar year (Jan-Dec).<br>For southern hemisphere: | 50 |
| 7 | Cummulative GPP | MOD17A1 | 0.05° | monthly | Sum of monthly GPP over 12 months up to the previous month | 50 |
| 8 | Population density | GHS GPW4 | 0.05° | 5 yearly | Population density, transformed as $\log(1+x)$, interpolated to monthly from 5-yearly data | 51 |
| 9 | Road network density | GRIP4 | 0.25° | Snapshot | transformed as $\log(1+x)$ | 52 |
| 10 | Vegetation type fractions | MOD12Q1 / MCD12C1 | 0.0083° | Snapshot / yearly | Converted to fractions as described in the main text | 53, 54 |
| 11 | Lightning frequency | WWLLN | 0.0083° | monthly climatology |  | 55 |
|  | **Calibration** |  |  |  |  |  |
| 11 | Burned area | GFED4.1s | 0.25° | monthly |  | 9 |

**SI-Figure 1.**
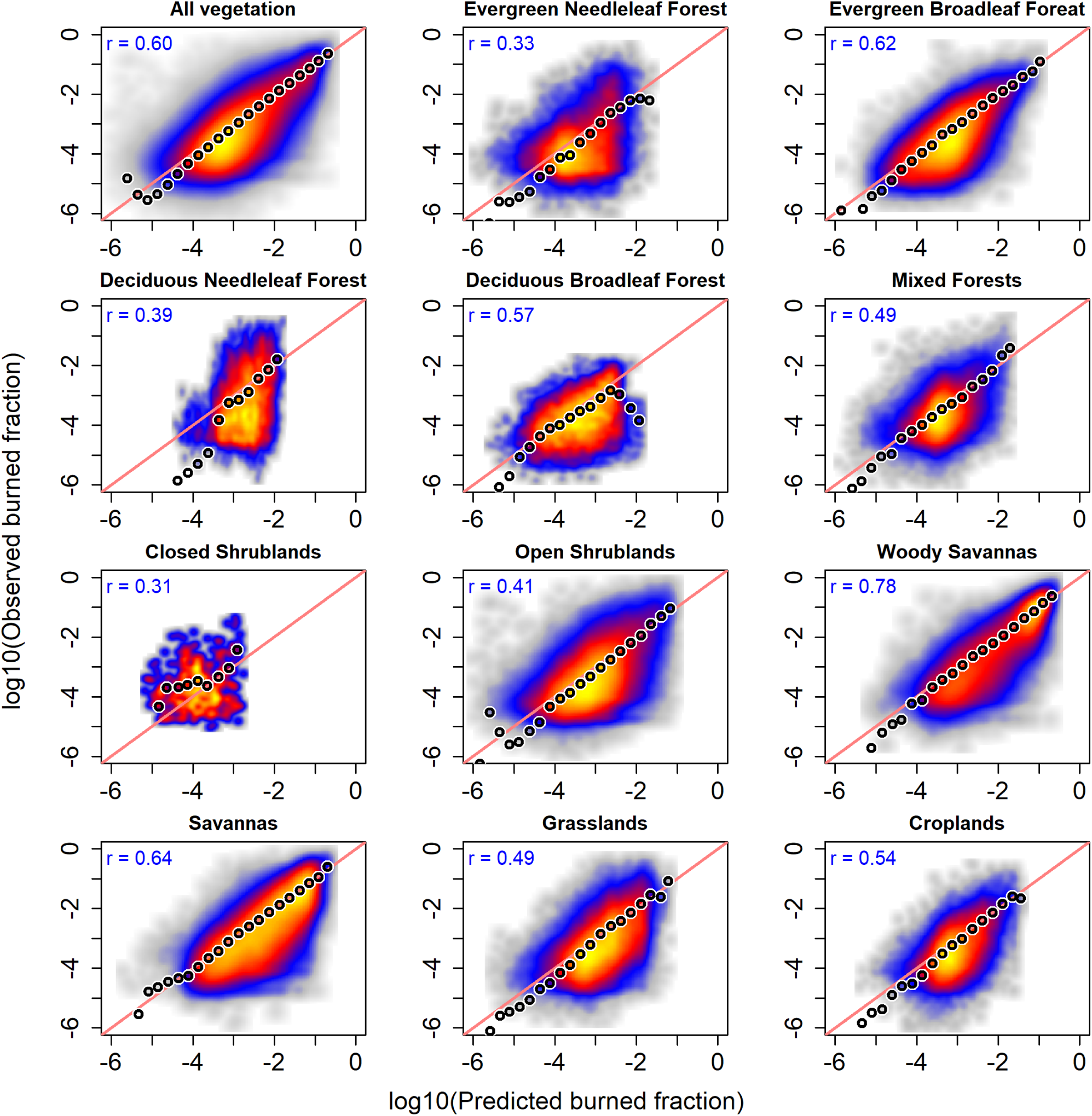
Observed vs predicted burned area for each forest type. Red line is the 1:1 line, and black/white circles show mean BA in each class. Density of points increases from grey to blue to red to yellow. Correlation is indicated in the top left corner. To identify the dominant PFT in each grid, we first excluded all grids with more than 50% non-vegetated or agricultural area (they were classified as non-vegetated and croplands respectively). Among the remaining grids, we ranked the types by abundance. If the most abundant type was at least 10% more abundant than the second most abundant one, we classified the grid as dominated by that type, or else as mixed vegetation.

**SI-Table 3.** Models used for sensitivity analysis, in the case where the best regional model from Table 1 does not include temperature. These models may have a substantial performance loss (Score difference > 5) compared to the best model.

| Region | T | IA | S | BA | LT | IA (det) | Score | Drop | Vars | Variables |
| --- | --- | --- | --- | --- | --- | --- | --- | --- | --- | --- |
| NHAF | 0.93 | 0.63 | 0.92 | 151.6 | -1.02 | 0.37 | 82.7 | -5.33 | 4 | pr, ts, cld, pop |
| CEAS | 0.71 | 0.59 | 0.79 | 15.4 | -0.07 | 0.57 | 82.2 | -3.34 | 5 | gppl1, pr, ts, cld, vp |
| EQAS | 0.82 | 0.92 | 0.83 | 2.1 | 0.03 | 0.94 | 94.4 | -0.54 | 4 | gppm1, pr, ts, cld |
| EUME | 0.81 | 0.29 | 0.65 | 2.2 | 0.02 | 0.34 | 70.5 | -0.19 | 2 | ts, cld |

**SI-Table 4.** Fire drivers in different regions as per our study and key previous studies.

|  | This study | Abatzoglou et al.<br>2018<sup>29</sup> | Andela et al.<br>2014<sup>42</sup> | Ponomarev et al.<br>2016<sup>56</sup> | Werf et al.<br>2008<sup>57</sup> | Aldersley et al.<br>2011<sup>31</sup> | Archibald et al.<br>2009<sup>30</sup> |
| --- | --- | --- | --- | --- | --- | --- | --- |
| BONA | moisture, fuel | aridity, climate |  |  |  |  |  |
| TCAM | climate, moisture, fuel | aridity |  |  |  |  |  |
| SA | climate, moisture, fuel | aridity |  |  | fdp |  |  |
| EUME | moisture |  |  |  |  |  |  |
| NHAF | moisture, pop |  | pr, pop, crop |  | no fuel, no aridity, | pop, wet, temp |  |
| SHAF | climate, fuel | fuel, climate | pr |  | no fuel, no aridity, | wet, crop | treecover, rainfall, dryseason, grazing |
| BOAS | climate, moisture, fuel | aridity, climate |  | aridity, temp |  | wet, tmp, crop, precip |  |
| CEAS | climate, fuel |  |  |  |  |  |  |
| SEAS | climate, moisture, fuel, pop | aridity |  |  |  |  |  |
| EQAS | moisture | fuel, climate |  |  |  |  |  |
| AUS | moisture, fuel | aridity, fuel, climate |  |  | npp | tree, precip |  |

**SI-Figure 2.**
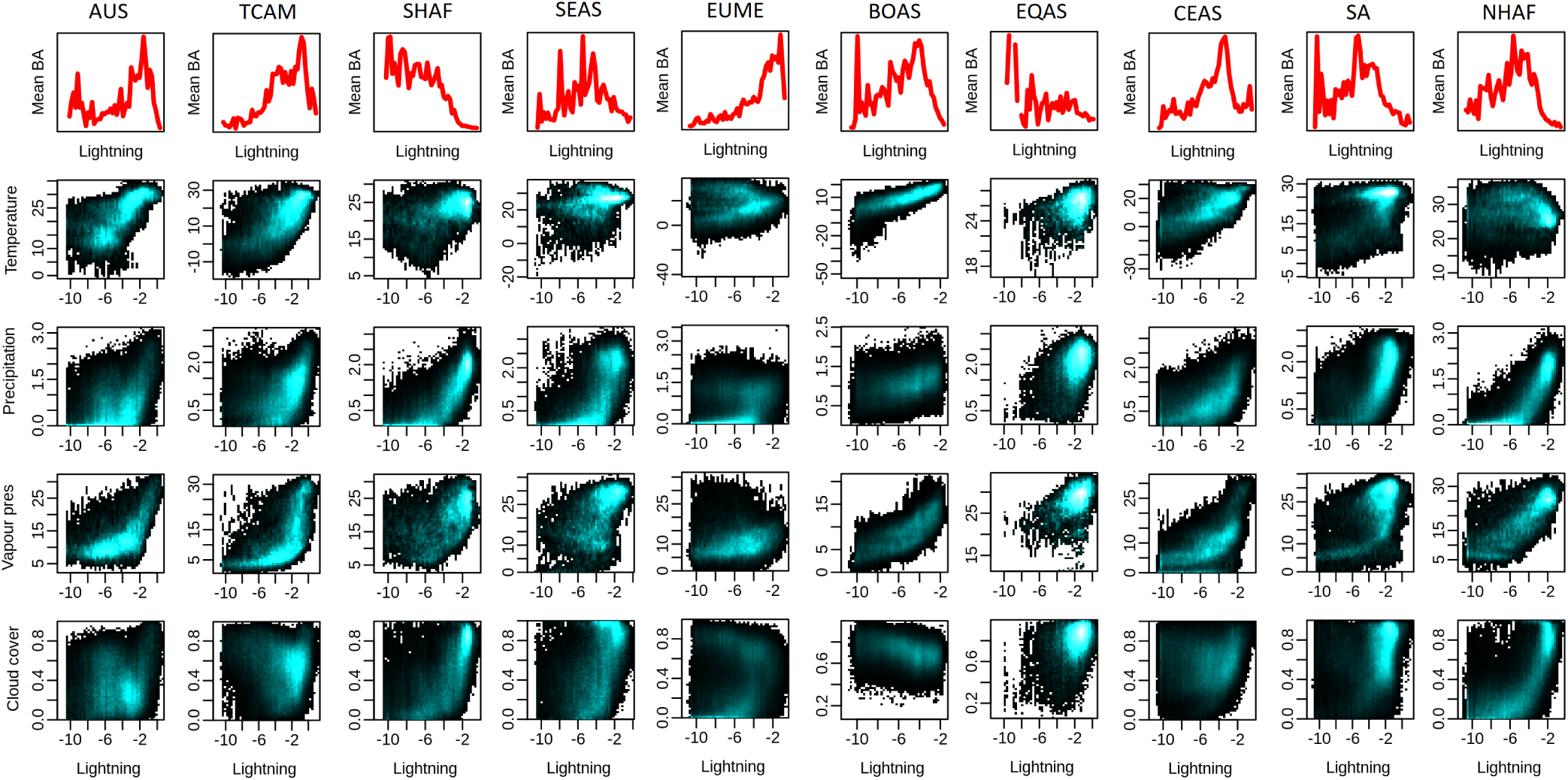
Relationships between lightning and other drivers.

**SI-Figure 3. Global temperature sensitivity for all months (GIF uploaded separately)**

**SI-Figure 4.**
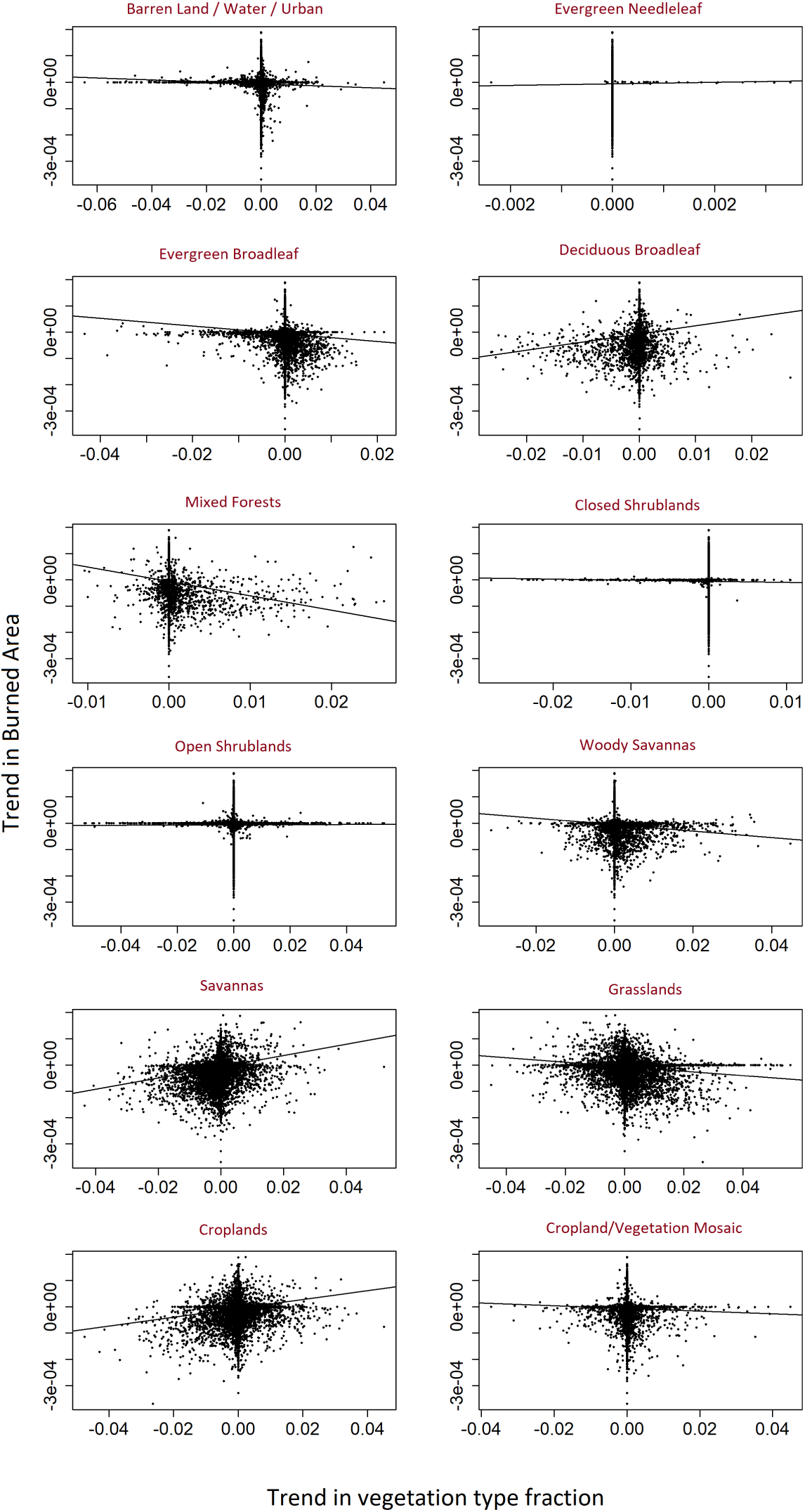
Correlation between trends in BA and trends in each vegetation type

**SI-Figure 5.**
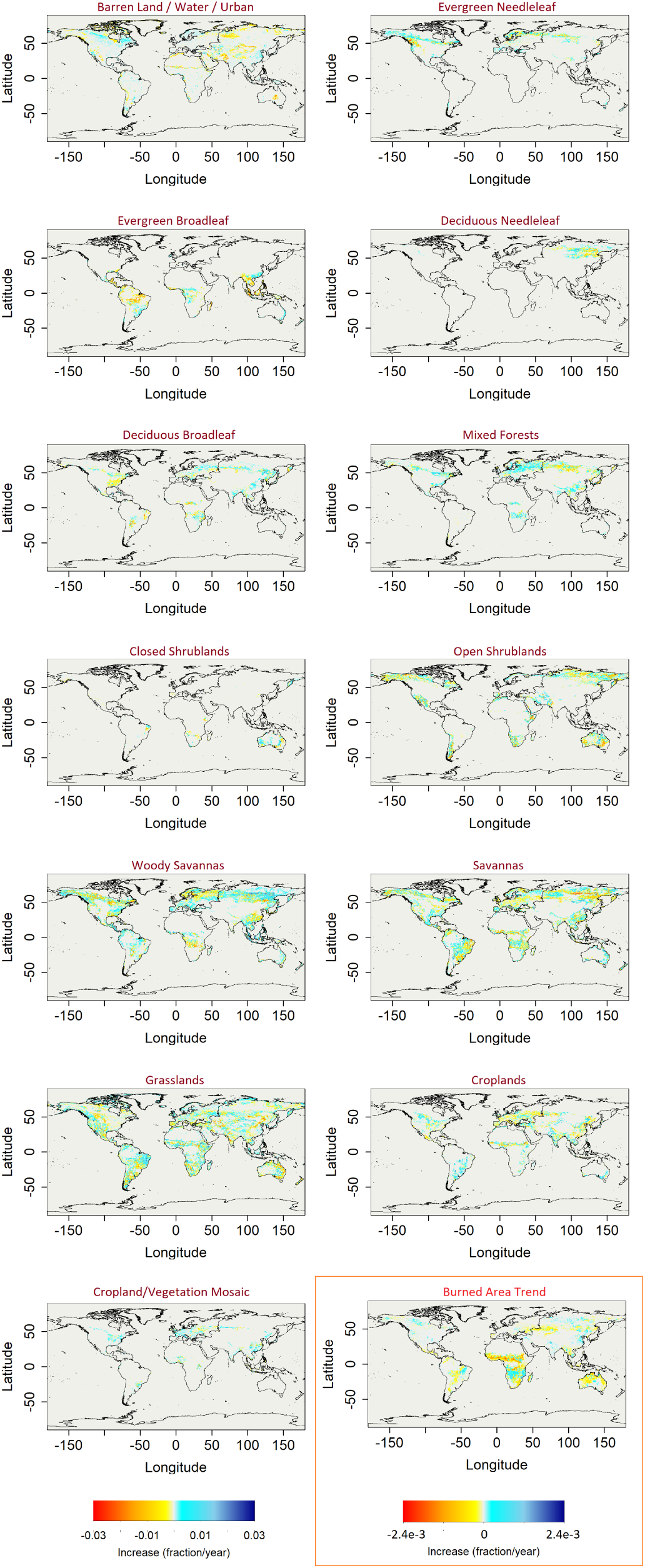
Trends in each vegetation type across the world from 2001-2017, along with the trend in burned area from 2001-2016. Also provided separately as a high-resolution file in SI.

**SI-Figure 6.**
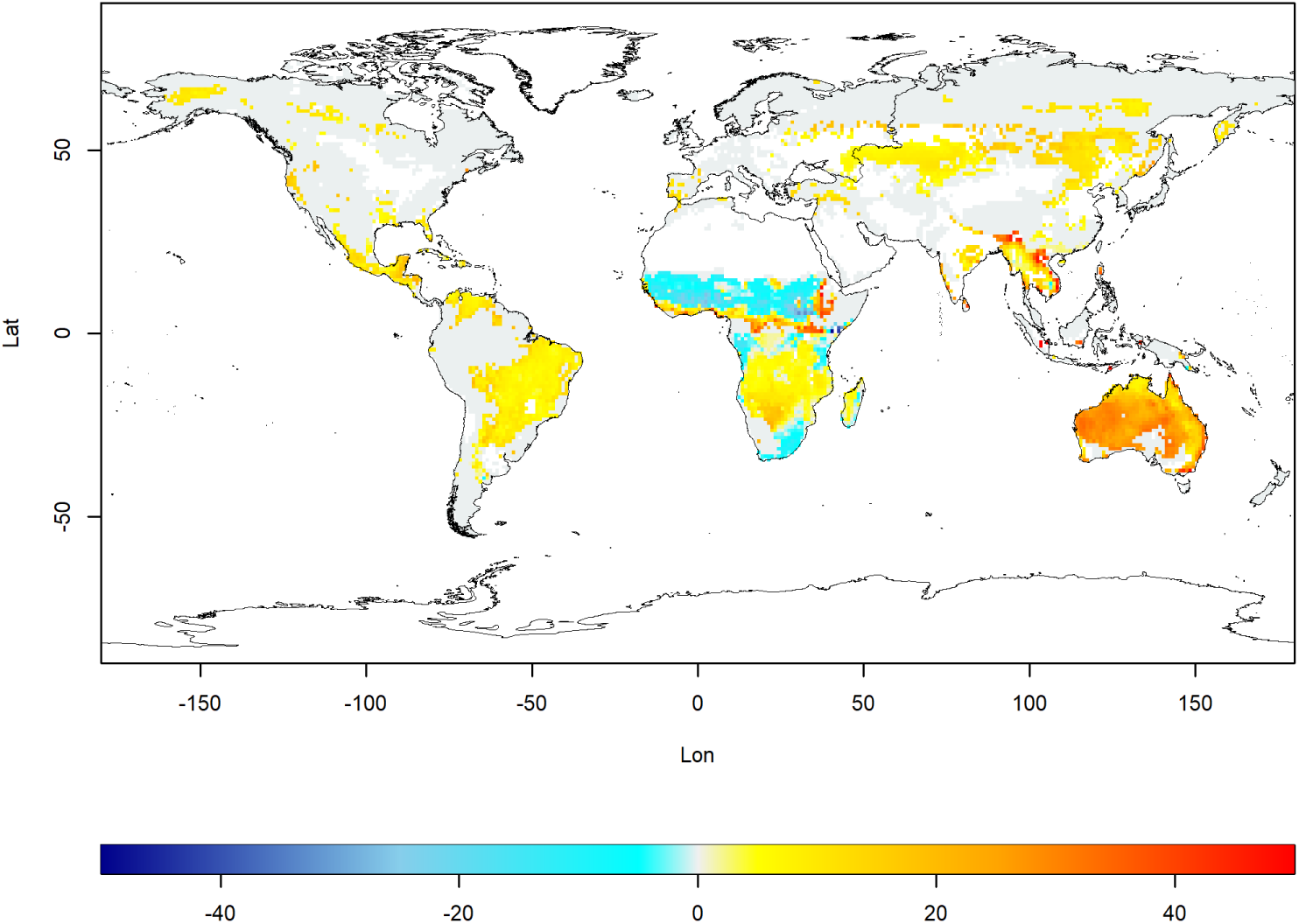
Sensitivity of fire to temperature (percent increase in burned fraction per unit change in temperature). Regions in eastern Himalaya and southeastern Australia stand out in terms of percentage increase in burned area fraction per unit rise in temperature. To avoid spurious values, we have removed cells with extremely low burned areas (< 1% of the cell burned). The projected high sensitivity of interior Australia is surprising given that temperatures are already high there, but this could be due to a lack of data prescribing a decline at very high temperatures (see discussion).

